# Virtual Parts Repository 2: Model-driven design of genetic regulatory circuits

**DOI:** 10.1101/2021.04.11.439316

**Authors:** Göksel Mısırlı, Bill Yang, Katherine James, Anil Wipat

## Abstract

Engineering genetic regulatory circuits is key to the creation of biological applications that are responsive to environmental changes. Computational models can assist in understanding especially large and complex circuits where manual analysis is infeasible, permitting a model-driven design process. However, there are still few tools that offer the ability to simulate the system under design. One of the reasons for this is the lack of accessible model repositories or libraries that cater for the modular composition of models of synthetic systems that do not yet exist in nature. Here, we present the Virtual Parts Repository 2, a resource to facilitate the model-driven design of genetic regulatory circuits, which provides reusable, modular and composable models. The repository is service-oriented and can be utilized by design tools in computational workflows. Designs provided in Synthetic Biology Open Language documents are used to derive system-scale and hierarchical Systems Biology Markup Language models. We also present a rule-based modeling abstraction based on reaction networks to facilitate scalable and modular modeling of complex and large designs. This modeling abstraction incorporates design patterns such as roadblocking, distributed deployment of genetic circuits using plasmids and cellular resource dependency. The computational resources and the modeling abstraction presented in this paper allow computational design tools to take advantage of computational simulations and ultimately help facilitate more predictable applications.

**Graphical TOC Entry:** 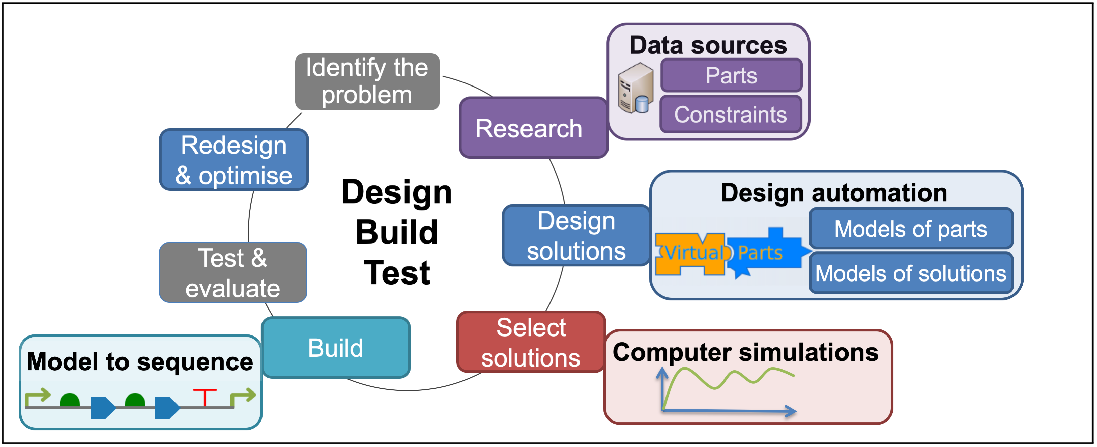

---

Synthetic biology is moving towards data-driven design applications (*1*). DNA fragments can be represented as electronic records ready to be composed virtually into designs for complex genetic circuits. Whether designs are created manually in a computer-aided environment or created computationally using heuristic approaches, these designs may need to be verified or optimized (*2*). The resulting systems can be exceedingly complex, depending on several parameters, and can show non-linearity between their inputs and outputs. It is almost impossible to predict a system’s resulting biological behavior intuitively as the number of biological components increases. Moreover, the use of components in different combinations and arrangements can give rise to a vast number of designs (*3*). However, not all designs are biologically viable.

Genetic design automation (GDA) has gained an interest to engineer biological systems with desired phenotypes (*4–8*). GDA tools typically address some of the design-build-test related tasks. As in other engineering disciplines (*9*), these tasks include identifying and researching the problem, designing alternative solutions, choosing from these alternatives, and fabricating and testing prototypes. One way to bridge the design and fabrication processes and to improve the predictability of genetic circuit designs is to apply model-driven design methodologies.

Model-driven design methodologies have already been proven to be a valuable tool to map a virtual design to a physical system for large-scale synthetic biology (*10, 11*). Computational models can capture the biochemical properties of biological parts. The aggregation of these models can then be used to understand the dynamics of a system built from individual parts (*12*).

Model-driven design frameworks should be easy to incorporate into existing tools via computational workflows; it is not practical or cost-effective to develop a single tool to accomplish complex tasks. Data standards are crucial to implement workflows when transforming the output from one tool into the input for another tool. Each tool can then be treated as a building block with inputs and outputs that are computationally interoperable with the other tools in a workflow (*13–15*).

The Systems Biology Markup Language (SBML) (*16*) is a widely used data standard for computational modeling of biological systems. In SBML models, *species* entities represent biological molecules such as proteins, DNA and signaling molecules. The *reaction* entities capture how these species interact, forming reaction networks. These reactions can be used to analyze how the incorporation of a biological part affects a reaction network and, hence, the overall behavior of a biological system. SBML Level 3 provides additional features to facilitate the hierarchical composition of computational models and their reuse as submodels (*17*). These submodels can be extended with *port* entities, which can be mapped to entities from different models.

The Synthetic Biology Open Language (*18–20*) (SBOL) is another data standard that is highly adopted in GDA. While SBML can be used to capture the dynamics of a genetic circuit, SBOL provides a machine-accessible definition of the circuit in terms of the circuit’s building blocks such as proteins, DNA-based components, and how these components are positioned. Different constraints, including molecular interactions, can be qualitatively represented. Moreover, designs can be defined hierarchically by reusing existing parts. In addition to the explicit data schema, applications can use custom metadata or annotations. This flexibility is due to the representation of SBOL documents as graphs, in which the interpretation of unknown nodes and edges can be left to applications.

SBOL and SBML are already increasingly being used together to capture genetic circuit designs and to predict part composability. Tools, such as iBioSim (*7*), Tellerium (*21*) and BMSS (*5*), provide a mapping between genetic designs and computational models using SBOL and SBML. In particular, the ability to store custom metadata in both SBOL and SBML (*22, 23*) allows the cross-referencing of design and modeling entities, and hence to provide conversions from designs to simulatable models (*14, 24, 25*) and vice versa (*22, 26*). As a graph language, SBOL can be directly stored in graph repositories. One such database framework is SynBioHub (*27*), which is used to store and share genetic designs. Design-related resources in a SynBioHub instance can be resolved using their uniform resource identifiers (URIs). It is desirable for model-driven design tools to access design information that is curated and stored in publicly available repositories.

Genetic regulatory circuits involve the production of gene products and the effect of these products on the activation and inhibition of genetic elements, such as inducible or repressible promoters. Ordinary differential equations (ODEs) and reaction networks can provide an adequate approximation to represent biological interactions and to analyze the resulting models (*28*). These mathematical equations are developed based on a modeling abstraction, which often does not provide a one-to-one mapping between biochemical reactions and corresponding modeling entities (*29*). It is desirable to develop modeling abstractions that accommodate scalability when models are composed computationally.

Modeling abstractions should also be compatible with the deployment scenarios of genetic regulatory circuits. Design patterns based on the distribution of genetic circuits have already emerged (*30, 31*). A key feature to control cellular behavior is varying copy numbers to tune the input-output behavior of regulatory networks (*32*). Deployment scenarios may involve chromosomal integration or using plasmids. The former can be applied to guarantee a single copy of a genetic circuit and is commonly applied to transform *Bacillus subtilis* cells (*33*). However, a more common approach for *Escherichia coli* is to deploy using low, medium or high copy plasmids of varying numbers (*34, 35*). A genetic circuit can then be split into sections, which can be deployed using different plasmids (*36*).

Roadblocking is another design pattern to implement biological logic gates (*6, 30, 34*). Roadblocking occurs when a transcription factor (TF) prevents the movement of RNA polymerase (RNAP) and stops downstream transcription (*37*). Flexible modeling approaches are necessary to represent this biological concept and to design reusable logic gates.

We previously developed the Virtual Parts Repository (*38*) version 1 (VPR1), which provides reusable and modular models of biological parts and interactions (*39*). These models are called virtual parts and can be used to create system-scale models computationally (*40*). VPR1 has a built-in relational repository, which enables the querying of molecular constraints between different parts. This data warehousing approach requires that all the required information is collected, transformed and integrated using the VPR1 data schema. VPR1 can store definitions of simple parts, although hierarchical design information cannot be stored and retrieved. Exporting information about biological descriptions of parts is limited. VPR1 uses SBML Level 2 to represent virtual parts and the resulting models. As a result, the model composition process relies on model annotations to connect inputs and outputs of different models.

This paper presents the second version of the Virtual Parts Repository (VPR2). VPR2 has been developed using a modular architecture including components for a Web-based repository, a web service, a client library for computational tools, and a standalone data library to retrieve data from remote SBOL repositories (Figure 1). The web service can be used to retrieve virtual parts and to create computational models of genetic circuits. VPR2 has a graph-based repository and allows browsing of the underlying data. It has also been developed as a modular service to take advantage of community-driven data repositories, such as a SynBioHub instance. The composition of models is facilitated via hierarchical SBML Level 3 models that rely on submodels. The resulting models can also be incorporated into other models. Importantly, VPR2 uses SBOL as a domain-specific language to control the composition of models and to specify design-related constraints. Tools can submit detailed genetic circuit descriptions, or the order and types of genetic parts to retrieve the rest of the information from a remote data repository. This service-oriented VPR2 is ideal for complex and computational workflows that utilize data standards.

**Figure 1:**
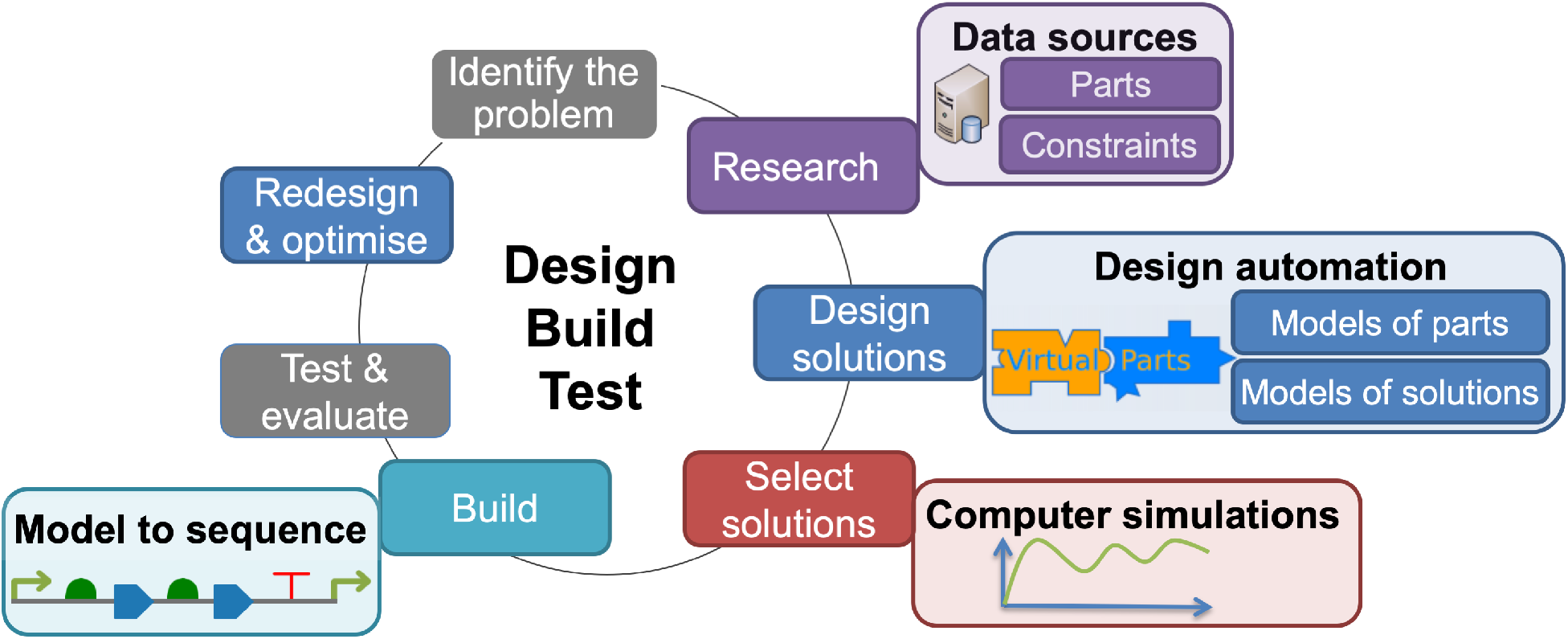
The VPR2 overview. VPR2 provides a data layer for tools to access information about parts and design constraints. This information is used to derive modular, reusable and composable models. The resulting system-scale models are used to identify biological solutions via simulations.

We also present a new modular modeling abstraction that allows flexible representation of biological systems using reaction networks. Complex interactions are represented using simple rules. This abstraction simplifies the modeling of genetic circuits and is scalable as the number of parts increases. Another new feature of the VPR2 modeling approach is incorporating the roadblocking concept. Any number of promoters or binding sequences can be included in a circuit in any order. Contextual information is incorporated into models via sigma factors, which can play significant roles in regulating transcription and controlling different cellular states (*41*). Such data can help design genetic circuits in changing environments and synchronize cellular resources. Moreover, this modeling abstraction allows the splitting of genetic circuits into subcircuits, each of which can be defined with different copy numbers.

## 1 Results and discussion

The Virtual Parts Repository version 2 (VPR2) has been developed to facilitate GDA by providing a computational framework to construct models of desired systems that can be tested via computer simulations (Figure 1). These simulations provide insights into the temporal behavior of genetic circuits composed of different parts. The model construction process can be automated to search large biological design spaces to find alternative solutions, each of which is represented as a single system-scale model. Models satisfying the requirements can then be used to derive genotypes that encode desired phenotypes.

### Computational modeling and data integration as a service

VPR2 provides a computational mechanism to access information about parts and constraints to derive models of parts and to create system-scale models. This framework is service-oriented and has a modular architecture (Figure 2). VPR2 comes with a built-in Resource Description Framework (RDF) graph repository, which can natively store SBOL documents. In addition to the genetic descriptions of parts and complex circuits, VPR2 related metadata can be embedded within SBOL entities to be stored in the repository. Data are queried using a graph pattern language (*42*). Consequently, VPR2 queries are generic and can be executed over any RDF graph repository with SBOL data. Using this approach, VPR2 has been designed to work with SynBioHub repositories, which store native SBOL data and provide HTTP endpoints for standard RDF queries.

**Figure 2:**
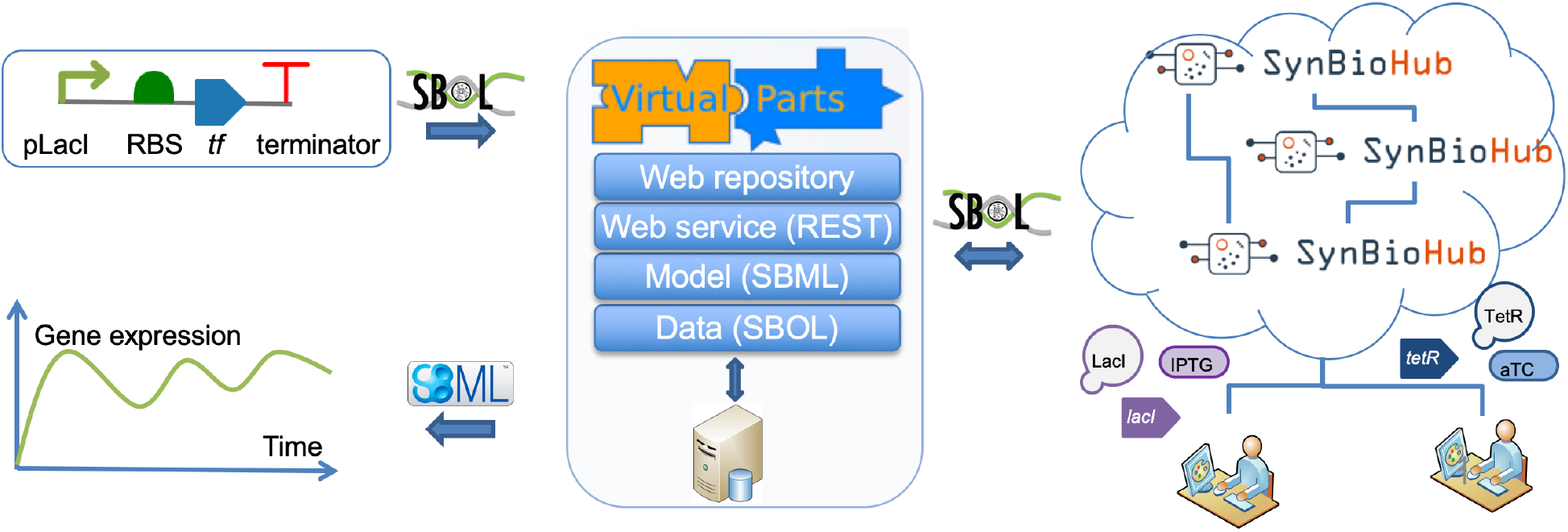
The VPR2 connected mode. Tools specify genetic circuits as ordered lists of parts using SBOL. VPR2 then searches for additional information about these parts via a specified design repository to create SBML models. SBOL designs are submitted to the VPR2 web service, which relies on model and data layers. The data layer uses SBOL to construct queries and to populate designs with detailed information. The modeling layer converts the resulting SBOL documents into SMBL models.

The primary data access method is provided by the VPR2 web service, which defines endpoints for model composition operations and model retrieval. The produced SBML models are hierarchical and may be formed of submodels, which import models of parts and interactions. A publicly available website has been developed for manual access. This web interface provides functionality to browse and search for models of biological parts and interactions from the VPR2 repository. In addition to using the web service, tools can use the VPR2 data library to return information about genetic parts, designs and interactions.

### Model-driven design using virtual parts

VPR2 relies on SBOL to specify genetic circuit designs and related constraints. Design specification information in these SBOL files is used to derive SBML models depicting the behaviors of desired systems. VPR2 can be used in *connected* or *disconnected* mode to derive SBML models or in *data* mode to enrich existing SBOL designs.

In the connected mode, tools provide basic design information and use VPR2 to retrieve quantitative and simulatable models. It is assumed that these tools do not have information about how different biological parts work together; SBOL is used to specify the types and order of parts. VPR2 then retrieves detailed information about these parts from a specified repository, which can be the local VPR2 repository or a remote SynBioHub instance. This detailed information is used to construct SBML models, which can be simulated via existing simulators such as COPASI (*43, 44*). Figure 2 depicts the use of the VPR2 framework in the connected mode. In this exemplar, a simple genetic circuit comprises a promoter, a ribosome binding site (RBS), a coding sequence (CDS) and a terminator. Consequently, VPR2 returns an SBML model that captures the circuit’s dynamic and temporal behavior. VPR2 provides direct access to design information by providing different search methods.

Tools can use these methods to initialize designs or to extend them. This information can be regarded as a biological network in which nodes represent parts and edges represents interactions. Hence, SBOL designs can be created by visiting different repository nodes representing parts and extending the designs based on the neighborhoods with other nodes. In the disconnected mode, tools provide all of the qualitative information required to create quantitative and simulatable SBML models. This information may include the order and types of biological parts and details about molecular interactions between these parts. Rate parameters can be provided in the form of custom annotations. These annotations are controlled using a set of VPR2 terms, and the values may indicate rate parameters. Figure 3 shows an example use of the VPR2’s disconnected mode. In the example design, the CDS part encodes a TF, which then transcriptionally inhibits the promoter. VPR2 returns the corresponding SBML model capturing the negative autoregulatory behavior of the genetic circuit.

**Figure 3:**
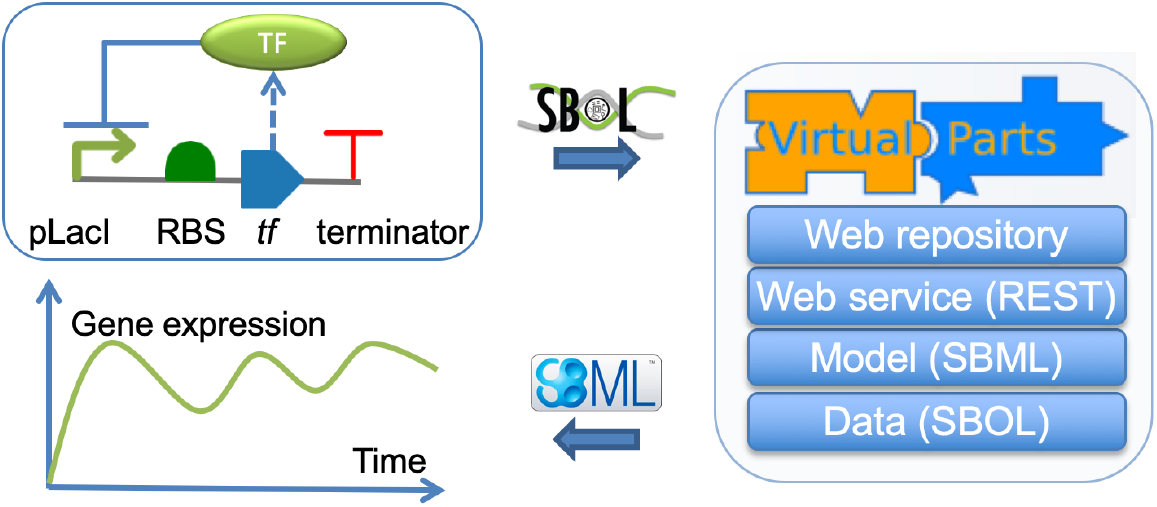
The VPR2 disconnected mode. No repository is involved in creating models. The disconnected mode acts as an SBOL-to-SBML converter. It is assumed that SBOL documents include all of the required information to derive SBML models.

VPR2 uses its modeling abstraction in both connected and disconnected modes. However, specialized tools in model-driven design technologies may depend on different modeling abstractions. VPR2’s *data* mode provides a data integration layer for such tools.

In the data mode, basic information about a genetic circuit in an SBOL document is used as input to retrieve more detailed information. Queries are directed to a specified repository to return information about molecular interactions and additional details about biological parts. Figure 4 shows the use of the data mode. Here, basic information about a genetic circuit describing the types and order of its parts is submitted to VPR2, which subsequently queries the specified repository to find genetic production and transcriptional repression interactions between the circuit’s parts and gene products.

**Figure 4:**
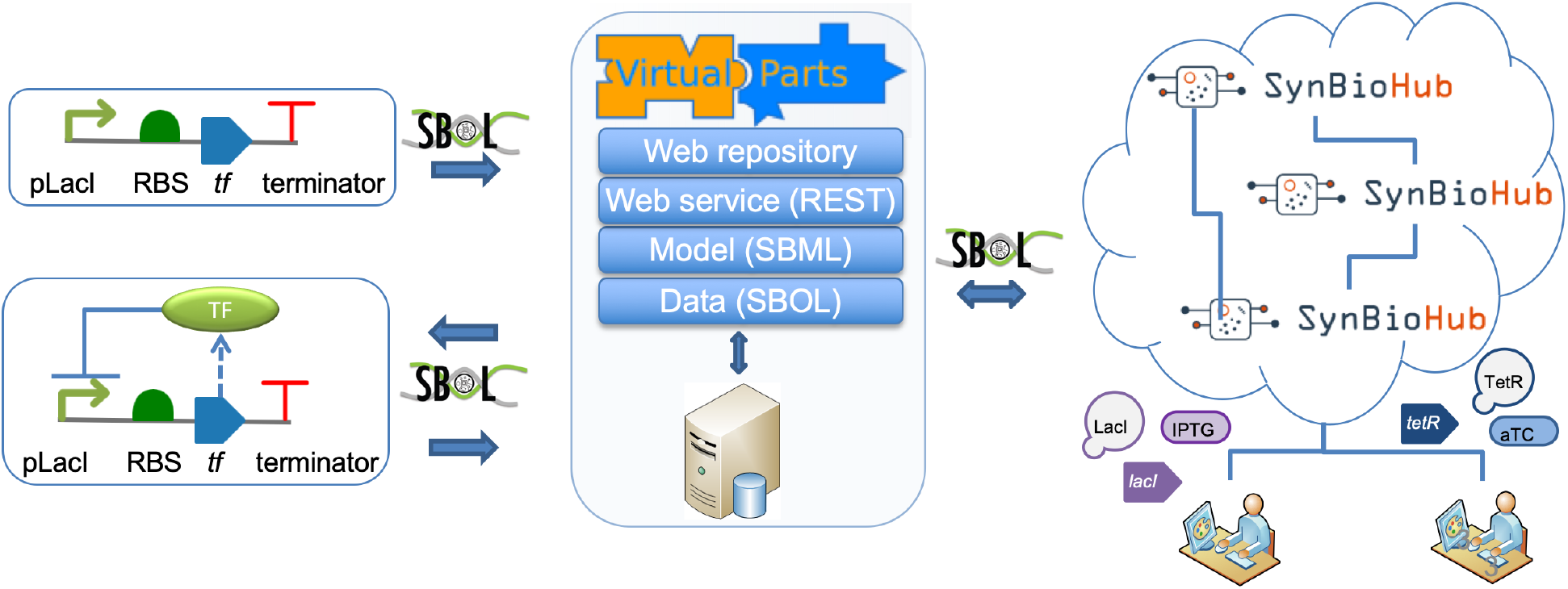
VPR2 *data* mode acts as the data integration layer. This mode is used to populate SBOL documents with additional information about parts and molecular interactions.

### Template-based model composition

VPR2 facilitates the model-driven design of large and complex genetic circuits by providing composable models. The composition of models is standardized by instantiating models of parts and interactions from well-defined templates based on the SBML’s hierarchical model composition package (*45*). These templates are fragments of models and may contain definitions of *species* and a set of *reaction* and *rule* SBML entities. The instantiation of these templates involves importing templates as submodels and parameterizing template-specific information. Templates are provided for promoter, RBS, CDS, operator and spacer genetic parts. Interaction templates include binding, degradation, phosphorylation, dephosphorylation, dimerization, DNA binding, and promoter activation and inhibition.

The composition process for system-scale models is carried out hierarchically, involving both the templates and the virtual parts. The resulting system-scale models are also sub-models that can be incorporated into other models. An example model composition process for a negative autoregulatory circuit is shown in Figure 5. The ‘*lacI* circuit’ model built from virtual parts includes information to simulate the system’s behavior. The topmost model shown in the figure is called the ‘System Model’ and is a placeholder to import the resulting submodel for simulations. In this circuit, the LacI TF inhibits the pLacI promoter, and hence its own production. VPR2 provides virtual parts for the pLacI promoter, the RBS and the *lacI* CDS. While the promoter virtual part captures the production of mRNAs, the RBS virtual part provides details about the initiation of protein translation from mRNAs using the ribosome per second (*46*) (RiPS) signal. The CDS model includes details about the genetic production of protein molecules from the RiPS output of the RBS virtual part. The VPR2 models can depend on multiple templates, which can further be derived from other templates. For example, the promoter template captures the dynamics of mRNA molecules and thus imports transcription and degradation templates as submodels (Figure 5).

**Figure 5:**
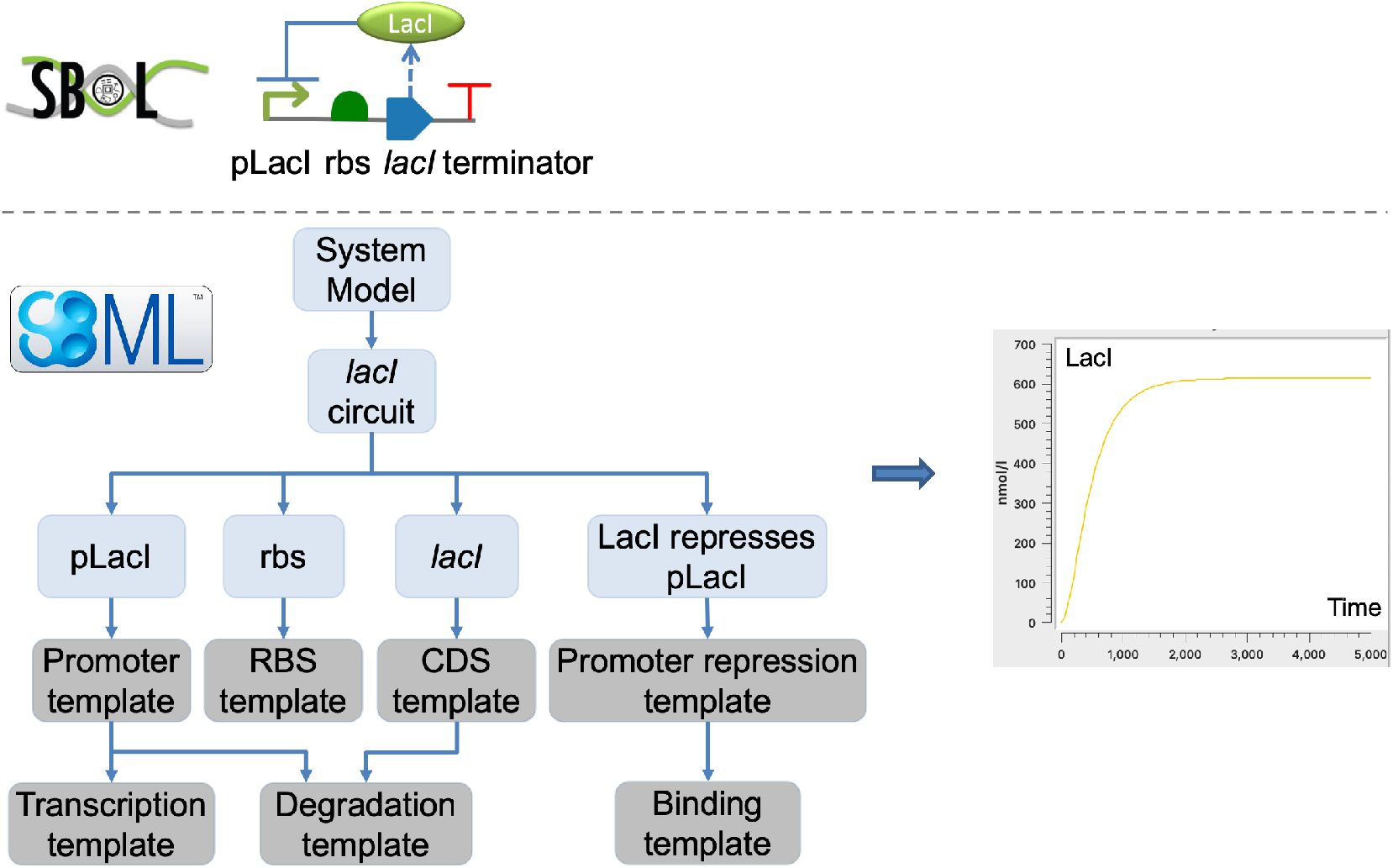
The VPR2 model composition. The LacI negative autoregulatory circuit is designed using a model-driven approach. The resulting ‘*lacI* circuit’ model is hierarchical and includes several other submodels.

The model composition process is facilitated via standard inputs and outputs, which can be defined for templates, virtual parts or system-scale models. VPR2 uses the SBML’s *port* entities to represent these inputs and outputs. The linking and replacement of these port entities provide a modular mechanism to compose SBML models. Firstly, these inputs and outputs are used to unify biological species’ meaning across different submodels. Secondly, they are used when instantiating templates to create and parameterize virtual parts by overwriting default parameters. Parameter overwriting can start from the topmost model and be passed to lower-level models. As default, parameters in lower-level models are used to calculate reaction fluxes, which then contribute to the concentration of molecules.

### A modeling abstraction for genetic design automation

This section presents a modeling abstraction that facilitates modular modeling of genetic regulatory circuits. This approach is scalable as the number of parts selected for a genetic circuit increases. The VPR2’s modeling abstraction is a state-based representation of biological molecules. States, such as a promoter bound or not bound to a TF, are represented as different species. The transitions between states are controlled via reactions representing the binding and unbinding of molecules. Additional reaction entities for different states of molecules are then used to provide simple rules rather than to use complex transfer functions to model the effect of multiple biological reactions. Transfer functions, such as Hill equations, are utilized to incorporate the impact of cellular resources.

In this modeling abstraction, promoters are represented as mRNA generators (Figure 6). An inducible promoter may have weak basal expression and may require an activator to reach its full potential (*47*). Conversely, a repressible promoter may be active until an inhibition signal is received via the binding of a repressor TF (*48*). Hence, in the VPR2’s modeling abstraction, mRNA production reactions are defined for promoters when they are in the unbound state (for constitutive, repressible and inducible promoters) or when they are bound to inducer TFs. Different rate parameters control how strong the corresponding production fluxes are. The reaction flux (*R*) for the production of mRNAs from a free or unbound promoter has the form:

**Figure 6:**
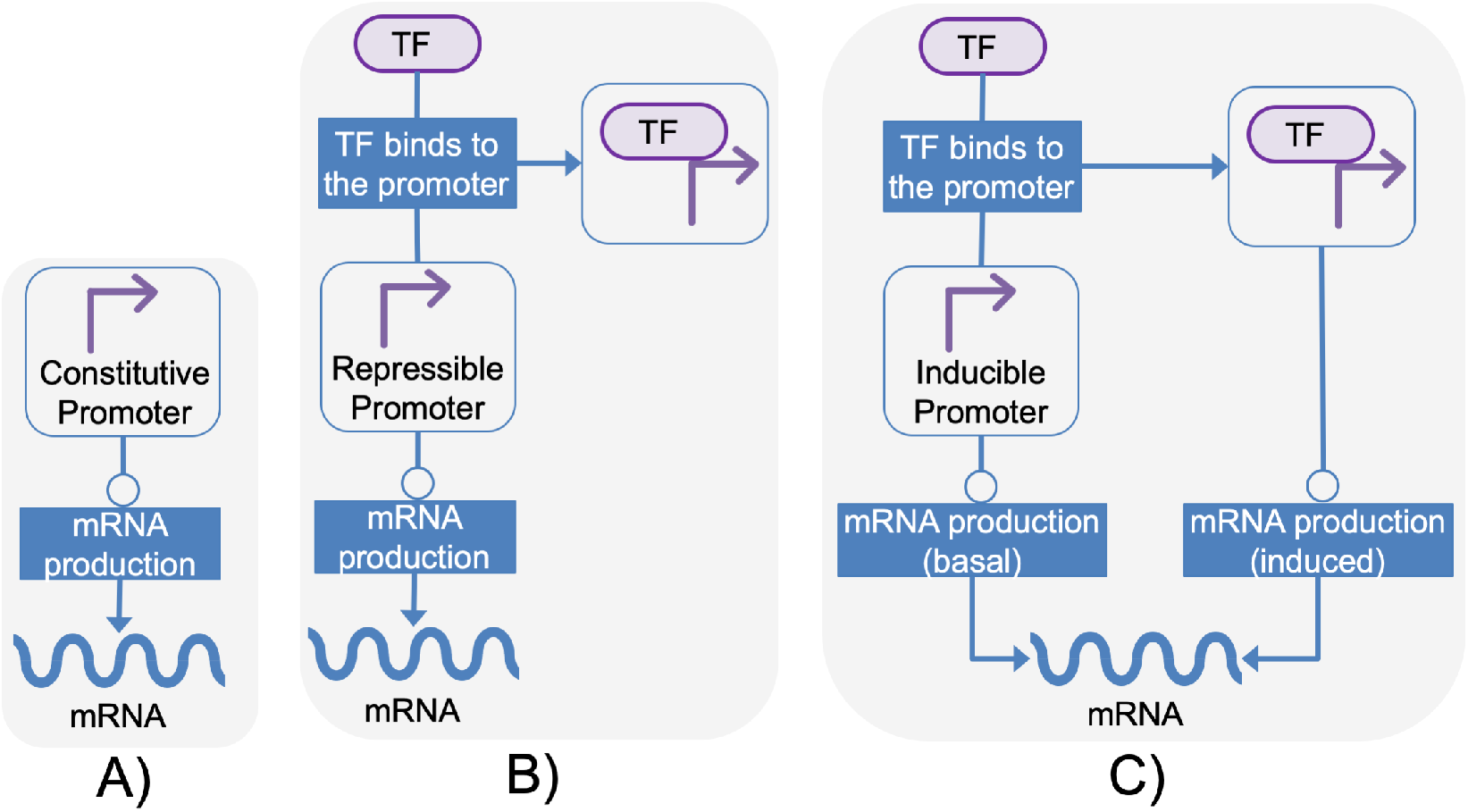
mRNA production from constitutive, repressible and inducible promoters. The diagram is created according to the Systems Biology Graphical Notation (*50*). Lines without arrows represent consumption, lines with arrows represent production, and lines with circles represent catalysis roles. A. A constitutive promoter. B. A repressible promoter. TF binds to the promoter and reduces the promoter’s probability of being free for mRNA production. C. An inducible promoter. When the promoter is free, mRNA is produced at a basal rate. When the promoter is bound to TF, it is activated.

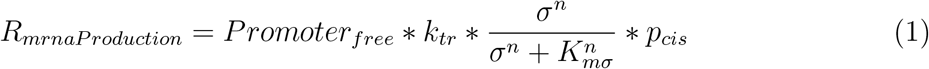

where *Promoter*_*free*_ is the number of free promoters, *k*_*tr*_ is the rate of transcription, and *σ* is the Sigma factor controlling the expression of the promoter. The sigma factor’s effect is described using 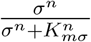, where *K*_*mσ*_ is the coefficient associated with the sigma factor, and *n* is the Hill equation (*49*). It is assumed that mRNA production is uniform for downstream genetic parts within a transcriptional unit. Therefore, the promoter activity is formulated as a function of the unbound downstream components to model RNAP elongation. Here, *p*_*cis*_ represents the probability of all downstream *cis* entities being free or unoccupied for a uniform mRNA transcription. *p*_*cis*_ is approximated as:

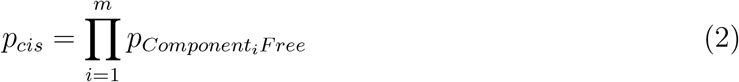

where *m* is the total number of genetic parts between a promoter and a downstream terminator, and 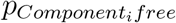 is the probability of the genetic part in position *i* being free. VPR2 allows for the roadblocking concept by incorporating the availability of each down-stream component 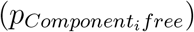, which can be approximated by dividing the number of unbound copies of a genetic part by the sum of its bound and unbound copy numbers (Equation 3). The following equation dynamically provides the probability of a genetic part being free or not based on the transient value of free copy numbers of that genetic part at time *t*.

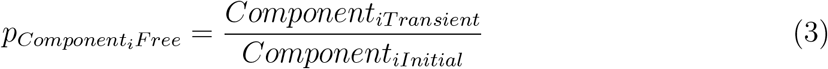

A repressible promoter uses the same formulation in Equation 1 for the mRNA production reaction flux (Figure 6B). Inhibition of the promoter is modeled implicitly by adding a reaction that represents the inhibitor’s binding to the promoter. This interaction affects the promoter’s probability of being free or not and takes away from the number of *Promoter*_*free*_ contributing to the accumulation of the promoter species in the bound state. This promoter occupation reaction flux can be represented as:

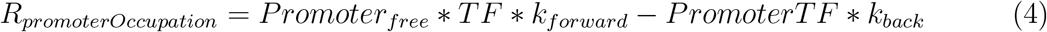

where *k*_*forward*_ is the forward reaction rate and *k*_*back*_ is the reverse reaction rate. In this reaction flux, both *Promoter*_*free*_ and *TF* are substrates, while *PromoterTF* is the product, which represents the complex formed by the promoter and the TF.

Regarding the activation of a promoter, promoter and TF complexes play an essential role by increasing the rate of RNAP binding to the promoter. The complex formation between the TF and the promoter is modeled using a similar approach to that used for repressible promoters (Figure 6C). However, when the promoter is bound to a TF, the promoter and the TF complex would also contribute to the mRNA production, in addition to the weaker basal expression from an unbound promoter. The mRNA expression for an activated promoter is modeled similarly as in Equation 1:

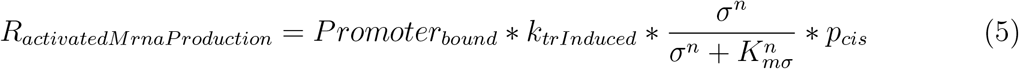

where *Promoter*_*bound*_ replaced *Promoter*_*free*_ to represent the promoter’s state when bound to an activator.

The RBS model converts mRNA signals into RiPS using the binding specificity between mRNAs and ribosomes (*39*). This specificity is a rate-limiting factor and can be affected by several factors, such as the Shine-Dalgarno sequence, the start codon and 5’ sequence, due to the formation of secondary structures (*51*). In VPR2’s modeling abstraction, a single rate parameter is used to capture these details. This parameter can be predicted using tools, such as RBS Calculator (*52*) and the UTR Designer (*53*). The RBS model is defined as:

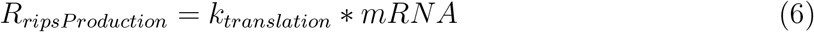

where *k*_*translation*_ is the rate of translation, and mRNA is the total number of mRNAs containing the RBS of interest.

The CDS model converts the RiPS reaction flux from an RBS model into an entity representing the gene product (*39*). The conversion is carried out using an ODE, which also incorporates the degradation of the gene product. A CDS virtual part provides entities to calculate the production flux and imports the degradation template, which is parameterized with a suitable degradation rate.

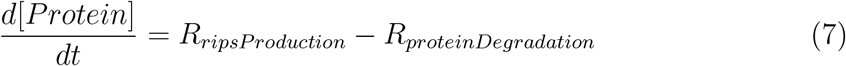

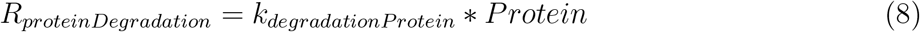

Other DNA-based components, such as operators, are represented as SBML species entities that can react as substrates. The inclusion of these components is modular. Whether these DNA-based components are bound or unbound changes the value of *p*_*cis*_ (Equation 1) to incorporate how mRNA transcription from preceding promoter parts is affected.

The interaction between an operator and a TF is modeled using the DNA binding template (Equation 9a). Another binding template is used when reactions involve modifiers or co-factors (Equation 9b). It is assumed that the concentration of such a molecule does not change. A specific form of binding template is defined for dimerization, where two molecules of the same type can form a complex (Equation 9c).

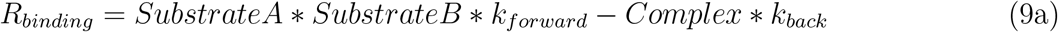

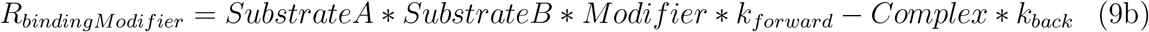

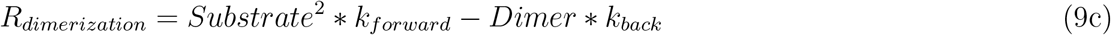

Templates also exist to model post-translational modifications (*38, 47*). The phosphorylation interaction is represented via a mechanism in which a phosphate donor transfers its phosphate to a phosphate acceptor. The reaction flux is shown in Equation 10a, where *Donor*~*p* and *Acceptor* are substrates, and *Acceptor*~*p* and *Donor* are products. An example of such an interaction is seen in bacterial two-component systems (*54*) formed of kinase and response regulator pairs. The phosphorylated form of a kinase protein can transfer its phosphate to a response regulator, which then becomes an active transcriptional regulator, for example, to induce gene expression. Kinases can be phosphorylated by external signals, acting as environmental sensors. Such environmental cues can be represented as modifiers, assuming that their overall concentration does not change, as shown in Equation 10b, where *Acceptor* is the substrate and its phosphorylated form *Acceptor*~*p* is the product.

For simplicity, an autodephosphorylation template is provided (Equation 10c).

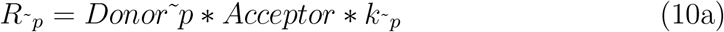

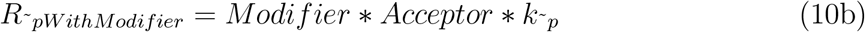

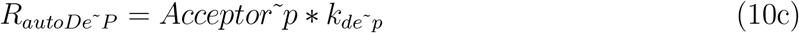

### An algorithm for genotype to phenotype mapping

VPR2 utilizes SBOL and SBML data standards to represent information relating to genotypes and corresponding phenotypes. Qualitative information defined in SBOL documents is used to derive SBML models. This conversion uses the following algorithm.

- Hierarchical genetic circuit designs are flattened into a list of transcriptional units in which genetic parts are explicitly ordered. Ordering is inferred using the exact start and end positions of genetic parts or using information about the relative arrangement of parts.
- A submodel is created for each design, a genetic part that does not have a parent design and acts as a container for child components.
- A single system-scale model is created connecting submodels of different designs.
- A submodel is also created for each transcriptional unit that is part of a design. It is assumed that mRNA production is uniform throughout each transcriptional unit. Sub-models for transcriptional units are then connected to submodels for designs.
- Each DNA-based genetic part is represented as a species within the transcriptional unit’s scope to which the genetic part belongs. These species connect submodels of transcriptional units with virtual parts and templates. For example, the same CDS with two copies would have two different species contributing to the corresponding gene product’s production. Similarly, multiple uses of an operator in the same or different designs would continue draining the number of TFs available due to binding and unbinding reactions dynamically.
- SBML species for non-DNA parts such as proteins and protein complexes are represented globally to connect system-scale models with submodels. For example, a single protein species at the system-scale model links all CDS species that encode the same protein and participate in translation reactions via different mRNA species.
- The roadblocking concept and the probability of genetic parts being free or not is used to link species representing DNA-based genetic parts in a transcriptional unit (Equation 2).
- An mRNA species is created for each promoter part in a transcriptional unit. These mRNAs represent genetic production between promoters and downstream terminators. Transcriptional units can include multiple promoters in any order.
- As a default, each species’ initial copy number is set to one for DNA-based genetic parts. Copy number is parameterized through genetic designs. If a copy number is provided as a parameter through an annotation, initial values for species representing DNA parts are set to that copy number.

### A toggle switch example

This section describes the VPR2 framework to generate a system-scale model of a genetic toggle switch. Toggle switches are modular genetic devices that can change cellular behavior between different states. Starting with Gardner’s (*55*) implementation of a genetic toggle switch, several studies have demonstrated the construction of toggle switches using different approaches (*6, 35, 56*). The example below is based upon the work of Lugagne and co-workers (*35*), who implemented a genetic toggle switch that can be controlled by a real-time feedback mechanism. The system’s outputs were produced as a series of transcriptional inhibition cascades. Individual cells were controlled using a microfluidic system. The system’s anhydrotetracycline (aTc) and isopropyl-*β*-D-thiogalactopyranoside (IPTG) inputs controlled RFP and GFP expressions, respectively. The aTc and IPTG input molecules were released into the system and were periodically washed out. Different concentrations of these inputs were used to analyze equilibrium states and instability.

Different states of this toggle switch could be reconstructed using the VPR2 framework. Figure 7A shows an adapted version of the circuit. The corresponding virtual parts were created based on the work of Nielsen and co-workers (*34*), who used aTc and IPTG molecules to implement genetic logic gates. These virtual parts were parameterized for reported mRNA copy numbers.

**Figure 7:**
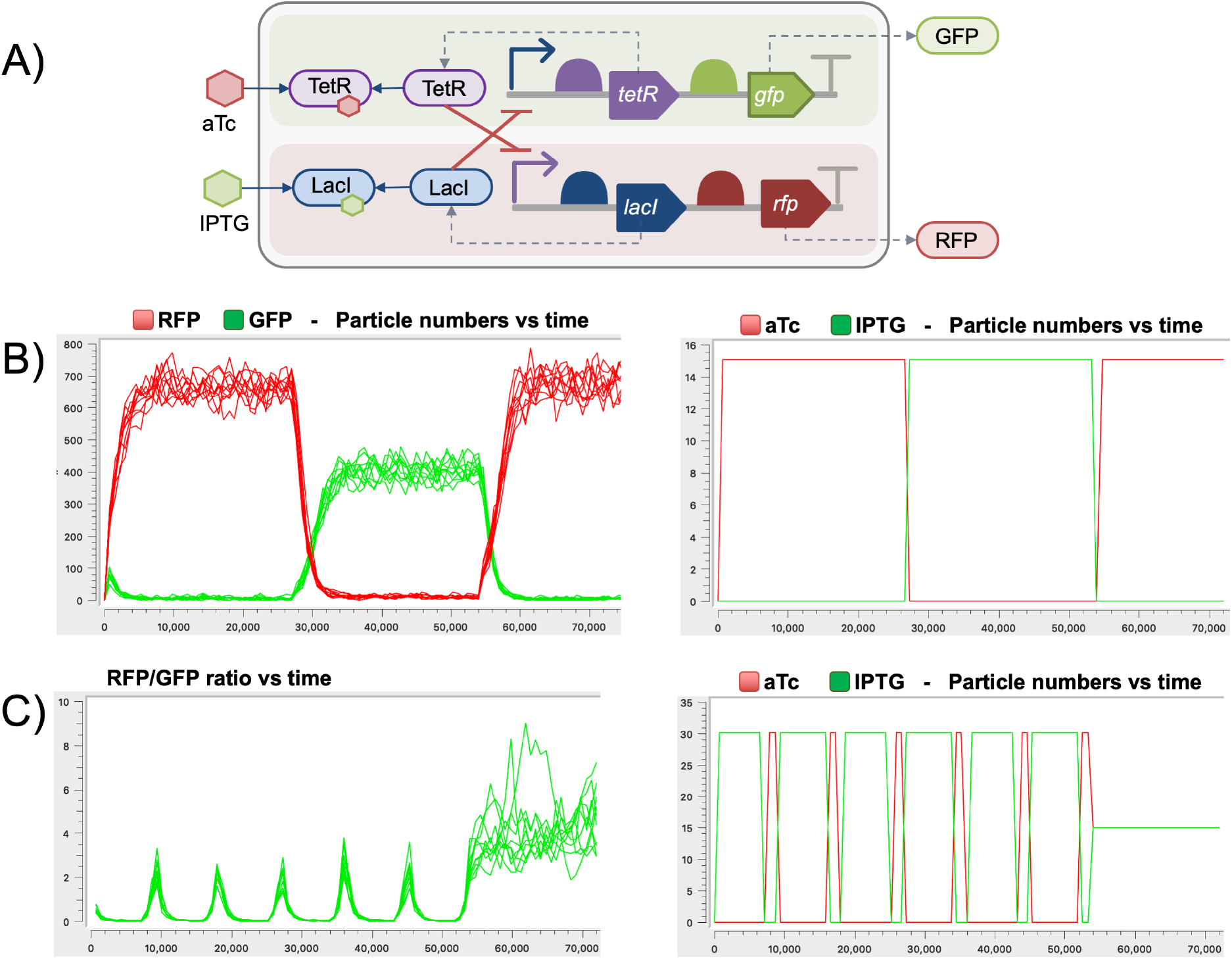
A toggle switch example, adapted from (*35*). X and Y axes represent time and particle numbers, respectively. A. The toggle switch genetic circuit design; GFP and RFP are produced in the presence of IPTG and aTc, respectively. B. GFP and RFP vs. time, and IPTG and aTc vs. time. The model was initially calibrated and tested against the work of Lugagne and co-workers (*35*). IPTG and aTc concentrations are varied every 7.5 hours. C. RFP/GFP ratio vs. time, and IPTG and aTc vs. time. The RFP/GFP ratio is shown when IPTG and aTc concentrations vary periodically (IPTG for 120 minutes and aTc for 30 minutes).

Similar to how Lugagne and co-workers initially tested this genetic circuit, a system-scale model was constructed to confirm relative RFP and GFP levels (Figure 7B). IPTG and aTc levels were then changed to control the RFP/GFP ratio and to verify that the system eventually favors RFP production (Figure 7C) (*35*). The models were simulated stochastically.

The VPR2 framework does not consider the diffusion of molecules. It is assumed that inputs such as IPTG and aTc have constant concentrations. Hence, these molecules are incorporated as modifiers. GFP and RFP production genetic circuits are assumed to be deployed using low copy plasmids (with ten copies).

### Discussion

VPR2 facilitates the modeling of genetic circuits and the automation of this process, providing reusable modeling and data services for computational tools. There is already an increasing number of tools that utilize computational simulations to predict the temporal behavior of genetic circuits. Developing computational models is not trivial. Models need to be carefully crafted every time, considering the biological constraints with a suitable mathematical framework.

Computational workflows can directly use the VPR2 framework. These workflows can either be implemented locally via a tool-specific task or distributed involving other tools. The framework is built on widely adopted SBOL and SBML standards to provide interoperability between different tools.

VPR2 decouples design and modeling processes. This separation allows focusing on design aspects and delegating the task of modeling to VPR2. Several design tools have already been developed using the SBOL standard (*57–59*). These SBOL compliant tools can take advantage of computational simulations by submitting designs of genetic regulatory circuits to VPR2 to retrieve simulatable SBML models.

Another modular approach that has been used to develop VPR2 is the decoupling of the design process and data storage. Although VPR2 comes with a local repository, other SBOL repositories can also be used. VPR2 can work with SynBioHub instances that are accessible via the Internet. Here, it is assumed that SBOL entities’ semantics are provided via recommended biological ontologies (*18, 25*).

In addition to providing a modeling service, VPR2 is also a data integration tool. DNA-based definitions of genetic circuits are enriched using information from SBOL repositories. This information is crucial when creating computational models. Hence, the VPR2’s integrative approach is also ideal for tools that can construct models. The VPR2’s data layer has already been integrated into iBioSim (*7*), a computational design and modeling tool. iBioSim provides SBOL designs to VPR2, which then enriches these designs. iBioSim then uses the enriched SBOL designs and converts them into SBML models using its modeling abstraction (*60*).

In addition to the computational resources, this paper presents a rule-based modeling abstraction using reaction networks. This modeling abstraction provides flexibility to capture the dynamics of genetic regulatory circuits built from individual parts. This approach is scalable and allows the integration of information as the complexity of genetic regulatory circuits increases. VPR2 takes the roadblocking concept into account, representing the implicit dependency between different genetic components. Moreover, VPR2 supports the modeling of distributed genetic circuits, where parts of these circuits can be deployed in different plasmids with different copy numbers (*36*).

The VPR2’s modular architecture allows the incorporation of new modeling abstractions and formats in the future. VPR2 assumes a uniform mRNA production and does not allow variations in genetic output from large operons. Moreover, terminators are assumed to be effective in stopping transcription. The modeling abstraction does not consider genetic parts in reverse directions. However, since VPR2 uses SBOL as a domain-specific language to specify modeling requirements, SBOL documents can be transformed to work with the VPR2 model generation process.

The computational resources presented in this paper facilitate the model-driven design of genetic regulatory circuits. VPR2 provides separation of design, modeling and data integration related tasks, and can be incorporated into computational workflows. Consequently, computational tools can take advantage of computational modeling and simulation, which is crucial to the design of predictable biological applications. Moreover, VPR2 provides a generic modeling abstraction to unlock the potential of designing genetic circuits and deriving computational models. This model-based genotype to phenotype mapping is especially important for genetic design automation in synthetic biology.

## 2 Methods

The VPR2 website has been developed using JavaServer Pages (JSP). The VPR2 web service is Representational State Transfer (REST) based and has been implemented using the Jersey framework. Its interface is available as a Web Application Description Language (WADL) file, a computationally accessible format providing information about the REST interface, parameters and return values of different endpoints. These endpoints can be used for particular operations, for example, to retrieve the model of a genetic circuit or a biological part. The web service can be accessed using HTTP POST operations. A Java client library has also been developed as a wrapper for the web service and provides programmatic access. The data layer has been developed in Java as a Maven project using libSBOLj (*61*) and can be used as a standalone library.

VPR2 is backed by an RDF repository. RDF4j was chosen for the default repository since it provides a simple approach to install and upload SBOL data. This repository currently includes the BacillOndex dataset (*62, 63*), which includes information about parts for *Bacillus subtilis*. Data retrieval from the local and other remote RDF repositories is facilitated using SPARQL (*64*), a graph pattern language for querying RDF-enabled graph repositories. These SPARQL queries are constructed using SBOL terms to unify accessing data.

Genetic regulatory designs are specified using SBOL. SBML Level 3 Version 1 is used to create hierarchical models. The simulation process requires that hierarchical models are flattened, and the inputs and outputs between submodels are explicitly mapped. VPR2 relies on SBML simulators to flatten these models. The COPASI (*44*) tool was used for simulations in our work. COPASI can import hierarchical SBML models and flatten them via libSBML (*65*).

### Availability

The VPR2 website is publicly available at http://www.virtualparts. org to manually browse data and computational models. The web service can be accessed computationally from http://www.virtualparts.org/virtualparts-ws/webapi. The VPR-data API and a client library to programmatically access the VPR web service are available as standalone Java libraries. Documentation about how to access and use these libraries, and different components of the VPR2 framework is available via the documentation section at http://www.virtualparts.org.

## References

1. Freemont, P. S. (2019) Synthetic biology industry: data-driven design is creating new opportunities in biotechnology. Emerg. Top. Life Sci. 3, 651–657.

2. Church, G. M., Elowitz, M. B., Smolke, C. D., Voigt, C. A., and Weiss, R. (2014) Realizing the potential of synthetic biology. Nat. Rev. Mol. Cell Biol. 15, 289–294.

3. Woodruff, L. B., Gorochowski, T. E., Roehner, N., Mikkelsen, T. S., Densmore, D., Gordon, D. B., Nicol, R., and Voigt, C. A. (2016) Registry in a tube: multiplexed pools of retrievable parts for genetic design space exploration. Nucleic Acids Res. 45, 1553–1565.

4. Appleton, E., Mehdipour, N., Daifuku, T., Briers, D., Haghighi, I., Moret, M., Chao, G., Wannier, T., Chiappino-Pepe, A., Belta, C., et al. (2019) Genetic Design Automation for Autonomous Formation of Multicellular Shapes from a Single Cell Progenitor. bioRxiv 807107.

5. Yeoh, J. W., Ng, K. B. I., Teh, A. Y., Zhang, J., Chee, W. K. D., and Poh, C. L. (2019) An Automated Biomodel Selection System (BMSS) for Gene Circuit Designs. ACS Synth. Biol. 8, 1484–1497.

6. Nguyen, T., Jones, T. S., Fontanarrosa, P., Mante, J. V., Zundel, Z., Densmore, D., and Myers, C. J. (2019) Design of Asynchronous Genetic Circuits. Proceedings of the IEEE 107, 1356–1368.

7. Watanabe, L., Nguyen, T., Zhang, M., Zundel, Z., Zhang, Z., Madsen, C., Roehner, N., and Myers, C. (2018) iBioSim 3: a tool for model-based genetic circuit design. ACS Synth. Biol. 8, 1560–1563.

8. Otero-Muras, I., and Banga, J. R. (2017) Automated design framework for synthetic biology exploiting pareto optimality. ACS Synth. Biol. 6, 1180–1193.

9. Haik, Y., and Shahin, T. Engineering Design Process, 2nd ed.; Cengage Learning, 2011.

10. Lenhard, J., and Carrier, M. Introduction: Mathematics as a Tool. In Mathematics as a Tool; Springer, 2017; pp 1–19.

11. Knuuttila, T., and Loettgers, A. Mathematization in Synthetic Biology: Analogies, Templates, and Fictions. In Mathematics as a Tool; Springer, 2017; pp 37–56.

12. Lux, M. W., Bramlett, B. W., Ball, D. A., and Peccoud, J. (2012) Genetic design automation: engineering fantasy or scientific renewal? Trends in Biotechnology 30, 120 –126.

13. Misirli, G., Madsen, C., de Murieta, I. S., Bultelle, M., Flanagan, K., Pocock, M., Hallinan, J., McLaughlin, J. A., Clark-Casey, J., Lyne, M., et al. (2017) Constructing synthetic biology workflows in the cloud. Eng. Biol. 1, 61–65.

14. Goni-Moreno, A., Carcajona, M., Kim, J., Martínez-García, E., Amos, M., and de Lorenzo, V. (2016) An implementation-focused bio/algorithmic workflow for synthetic biology. ACS Synth. Biol. 5, 1127–1135.

15. Myers, C. J., Beal, J., Gorochowski, T. E., Kuwahara, H., Madsen, C., McLaughlin, J. A., Misirli, G., Nguyen, T., Oberortner, E., Samineni, M., et al. (2017) A standard-enabled workflow for synthetic biology. Biochem. Soc. Trans. 45, 793–803.

16. Hucka, M., Finney, A., Sauro, H. M., Bolouri, H., Doyle, J. C., Kitano, H., Arkin, A. P., Bornstein, B. J., Bray, D., Cornish-Bowden, A., et al. (2003) The systems biology markup language (SBML): a medium for representation and exchange of biochemical network models. Bioinformatics 19, 524–531.

17. Hucka, M., Bergmann, F. T., Hoops, S., Keating, S. M., Sahle, S., Schaff, J. C., Smith, L. P., and Wilkinson, D. J. (2015) The Systems Biology Markup Language (SBML): language specification for level 3 version 1 core. J. Integr. Bioinform 12, 382–549.

18. Cox, R. S., Madsen, C., McLaughlin, J. A., Nguyen, T., Roehner, N., Bartley, B., Beal, J., Bissell, M., Choi, K., Clancy, K., Grünberg, R., Macklin, C., Misirli, G., Oberortner, E., Pocock, M., Samineni, M., Zhang, M., Zhang, Z., Zundel, Z., Gennari, J. H., Myers, C., Sauro, H., and Wipat, A. (2018) Synthetic biology open language (SBOL) version 2.2.0. J. Integr. Bioinform. 15, 20180001.

19. Galdzicki, M., Clancy, K. P., Oberortner, E., Pocock, M., Quinn, J. Y., Rodriguez, C. A., Roehner, N., Wilson, M. L., Adam, L., Anderson, J. C., Bartley, B. A., Beal, J., Chandran, D., Chen, J., Densmore, D., Endy, D., Grunberg, R., Hallinan, J., Hillson, N. J., Johnson, J. D., Kuchinsky, A., Lux, M., Misirli, G., Peccoud, J., Plahar, H. A., Sirin, E., Stan, G.-B., Villalobos, A., Wipat, A., Gennari, J. H., Myers, C. J., and Sauro, H. M. (2014) The Synthetic Biology Open Language (SBOL) provides a community standard for communicating designs in synthetic biology. Nat. Biotechnol. 32, 545–550.

20. Roehner, N., Beal, J., Clancy, K., Bartley, B., Misirli, G., Gruenberg, R., Oberortner, E., Pocock, M., Bissell, M., Madsen, C., Nguyen, T., Zhang, M., Zhang, Z., Zundel, Z., Densmore, D., Gennari, J., Wipat, A., Sauro, H., and Myers, C. J. (2016) Sharing Structure and Function in Biological Design with SBOL 2.0. ACS Synth. Biol. 5, 498–506.

21. Choi, K., Medley, J. K., König, M., Stocking, K., Smith, L., Gu, S., and Sauro, H. M. (2018) Tellurium: An extensible Python-based modeling environment for systems and synthetic biology. Biosystems 171, 74–79.

22. Misirli, G., Hallinan, J. S., Yu, T., Lawson, J. R., Wimalaratne, S. M., Cooling, M. T., and Wipat, A. (2011) Model annotation for synthetic biology: automating model to nucleotide sequence conversion. Bioinformatics 27, 973–979.

23. Neal, M. L., König, M., Nickerson, D., Misirli, G., Kalbasi, R., Dräger, A., Atalag, K., Chelliah, V., Cooling, M. T., Cook, D. L., et al. (2018) Harmonizing semantic annotations for computational models in biology. Brief. Bioinform. 20, 540–550.

24. Roehner, N., Zhang, Z., Nguyen, T., and Myers, C. J. (2015) Generating systems biology markup language models from the synthetic biology open language. ACS Synth. Biol. 4, 873–879.

25. Misirli, G., Nguyen, T., McLaughlin, J. A., Vaidyanathan, P., Jones, T. S., Densmore, D., Myers, C., and Wipat, A. (2019) A computational workflow for the automated generation of models of genetic designs. ACS Synth. Biol. 8, 1548–1559.

26. Nguyen, T., Roehner, N., Zundel, Z., and Myers, C. J. (2016) A converter from the systems biology markup language to the synthetic biology open language. ACS Synth. Biol. 5, 479–486.

27. McLaughlin, J. A., Myers, C. J., Zundel, Z., Misirli, G., Zhang, M., Ofiteru, I. D., Goñi-Moreno, A., and Wipat, A. (2018) SynBioHub: A Standards-Enabled Design Repository for Synthetic Biology. ACS Synth. Biol. 7, 682–688.

28. Karlebach, G., and Shamir, R. (2008) Modelling and analysis of gene regulatory networks. Nat. Rev. Mol. Cell Biol. 9, 770.

29. Alon, U. (2007) Network motifs: theory and experimental approaches. Nat. Rev. Genet. 8, 450–461.

30. Tamsir, A., Tabor, J. J., and Voigt, C. A. (2011) Robust multicellular computing using genetically encoded NOR gates and chemical ‘wires’. Nature 469, 212.

31. Ausländer, D., Ausländer, S., Pierrat, X., Hellmann, L., Rachid, L., and Fussenegger, M. (2018) Programmable full-adder computations in communicating three-dimensional cell cultures. Nat. Methods 15, 57.

32. Bradley, R. W., Buck, M., and Wang, B. (2016) Tools and principles for microbial gene circuit engineering. J. Mol. Biol. 428, 862–888.

33. Sonenshein, A. L., Hoch, J. A., Losick, R., et al. (2002) Bacillus subtilis and its closest relatives: from genes to cells.

34. Nielsen, A. A., Der, B. S., Shin, J., Vaidyanathan, P., Paralanov, V., Strychalski, E. A., Ross, D., Densmore, D., and Voigt, C. A. (2016) Genetic circuit design automation. Science 352, aac7341.

35. Lugagne, J.-B., Carrillo, S. S., Kirch, M., Köhler, A., Batt, G., and Hersen, P. (2017) Balancing a genetic toggle switch by real-time feedback control and periodic forcing. Nat. Commun. 8, 1671.

36. Bonnerjee, D., Mukhopadhyay, S., and Bagh, S. (2019) Design, fabrication and device chemistry of a 3-input-3-output synthetic genetic combinatorial logic circuit with a 3 input AND gate in a single bacterial cell. Bioconjugate Chem. 30, 3013–3020.

37. Hao, N., Palmer, A. C., Dodd, I. B., and Shearwin, K. E. (2017) Directing traffic on DNA—How transcription factors relieve or induce transcriptional interference. Transcription 8, 120–125.

38. Misirli, G., Hallinan, J., and Wipat, A. (2014) Composable modular models for synthetic biology. ACM J. Emerg. Technol. Comput. Syst. 11, 22.

39. Cooling, M. T., Rouilly, V., Misirli, G., Lawson, J., Yu, T., Hallinan, J., and Wipat, A. (2010) Standard virtual biological parts: a repository of modular modeling components for synthetic biology. Bioinformatics 26, 925–931.

40. Hallinan, J., Gilfellon, O., Misirli, G., and Wipat, A. Tuning receiver characteristics in bacterial quorum communication: An evolutionary approach using standard virtual biological parts. 2014 IEEE Conference on Computational Intelligence in Bioinformatics and Computational Biology. 2014; pp 1–8.

41. Haldenwang, W. G. (1995) The sigma factors of Bacillus subtilis. Microbiol. Mol. Biol. Rev. 59, 1–30.

42. Shadbolt, N., Berners-Lee, T., and Hall, W. (2006) The semantic web revisited. IEEE Intell. Syst. 21, 96–101.

43. Hoops, S., Sahle, S., Gauges, R., Lee, C., Pahle, J., Simus, N., Singhal, M., Xu, L., Mendes, P., and Kummer, U. (2006) COPASI—a complex pathway simulator. Bioinformatics 22, 3067–3074.

44. Bergmann, F. T., Hoops, S., Klahn, B., Kummer, U., Mendes, P., Pahle, J., and Sahle, S. (2017) COPASI and its applications in biotechnology. Journal of biotechnology 261, 215–220.

45. Smith, L. P., Hucka, M., Hoops, S., Finney, A., Ginkel, M., Myers, C. J., Moraru, I., and Liebermeister, W. (2015) SBML level 3 package: Hierarchical model composition, version 1 release 3. J. Integr. Bioinform. 12, 603–659.

46. De Lorenzo, V., and Danchin, A. (2008) Synthetic biology: discovering new worlds and new words. EMBO Rep. 9, 822–827.

47. Mukherji, S., and Van Oudenaarden, A. (2009) Synthetic biology: understanding biological design from synthetic circuits. Nat. Rev. Genet. 10, 859.

48. Cox, R. S., Surette, M. G., and Elowitz, M. B. (2007) Programming gene expression with combinatorial promoters. Mol. Syst. Biol. 3, 145.

49. Burrill, D. R., and Silver, P. A. (2010) Making cellular memories. Cell 140, 13–18.

50. Le Novère, N. (2015) Quantitative and logic modelling of molecular and gene networks. Nat. Rev. Genet. 16, 146.

51. Reeve, B., Hargest, T., Gilbert, C., and Ellis, T. (2014) Predicting translation initiation rates for designing synthetic biology. Front. Bioeng. Biotechnol. 2, 1.

52. Salis, H. M. The ribosome binding site calculator. In Methods Enzymol.; Elsevier, 2011; Vol. 498; pp 9–42.

53. Seo, S. W., Yang, J.-S., Kim, I., Yang, J., Min, B. E., Kim, S., and Jung, G. Y. (2013) Predictive design of mRNA translation initiation region to control prokaryotic translation efficiency. Metab. Eng. 15, 67–74.

54. Fabret, C., Feher, V. A., and Hoch, J. A. (1999) Two-component signal transduction in Bacillus subtilis: how one organism sees its world. J. Bacteriol. 181, 1975–1983.

55. Gardner, T. S., Cantor, C. R., and Collins, J. J. (2000) Construction of a genetic toggle switch in Escherichia coli. Nature 403, 339.

56. Lebar, T., Bezeljak, U., Golob, A., Jerala, M., Kadunc, L., Pirš, B., Stražar, M., Vučko, D., Zupančič, U., Benčina, M., Forstnerič, V., Gaber, R., Lonzarić, J., Majerle, A., Oblak, A., Smole, A., and Jerala, R. (2014) A bistable genetic switch based on designable DNA-binding domains. Nat. Commun. 5, 5007.

57. Zhang, M., McLaughlin, J. A., Wipat, A., and Myers, C. J. (2017) SBOLDesigner 2: An intuitive tool for structural genetic design. ACS Synth. Biol. 6, 1150–1160.

58. Der, B. S., Glassey, E., Bartley, B. A., Enghuus, C., Goodman, D. B., Gordon, D. B., Voigt, C. A., and Gorochowski, T. E. (2016) DNAplotlib: programmable visualization of genetic designs and associated data. ACS Synth. Biol. 6, 1115–1119.

59. Bates, M., Lachoff, J., Meech, D., Zulkower, V., Moisy, A., Luo, Y., Tekotte, H., Franziska Scheitz, C. J., Khilari, R., Mazzoldi, F., et al. (2017) Genetic Constructor: An Online DNA Design Platform. ACS Synth. Biol. 6, 2362–2365.

60. Misirli, G., Nguyen, T., McLaughlin, J. A., Vaidyanathan, P., Jones, T. S., Densmore, D., Myers, C., and Wipat, A. (2019) A Computational Workflow for the Automated Generation of Models of Genetic Designs. ACS Synth. Biol. 8, 1548–1559.

61. Zhang, Z., Nguyen, T., Roehner, N., Misirli, G., Pocock, M., Oberortner, E., Samineni, M., Zundel, Z., Beal, J., Clancy, K., et al. (2015) libSBOLj 2.0: a java library to support SBOL 2.0. IEEE Life Sci. Lett. 1, 34–37.

62. Misirli, G., Wipat, A., Mullen, J., James, K., Pocock, M., Smith, W., Allenby, N., and Hallinan, J. S. (2013) BacillOndex: An integrated data resource for systems and synthetic biology. J. Integr, Bioinform. 10, 103–116.

63. Misirli, G., Hallinan, J., Pocock, M., Lord, P., McLaughlin, J. A., Sauro, H., and Wipat, A. (2016) Data integration and mining for synthetic biology design. ACS Synth. Biol. 5, 1086–1097.

64. Allemang, D., and Hendler, J. Semantic web for the working ontologist: effective modeling in RDFS and OWL; Elsevier, 2011.

65. Bornstein, B. J., Keating, S. M., Jouraku, A., and Hucka, M. (2008) LibSBML: an API library for SBML. Bioinformatics 24, 880–881.

